# Predicting targets and costs for feral-cat reduction on large islands using stochastic population models

**DOI:** 10.1101/2020.06.12.149393

**Authors:** Kathryn R. W. Venning, Frédérik Saltré, Corey J. A. Bradshaw

**Affiliations:** Global Ecology, College of Science and Engineering, GPO Box 2100, Flinders University, Adelaide, South Australia 5001, Australia

**Author notes:** Corresponding author: Global Ecology, College of Science and Engineering, Flinders University, GPO Box 2100, Adelaide, South Australia 5001, Australia. **Declaration of interest** NIL.

## Abstract

Feral cats are some of the most destructive invasive predators worldwide, particularly in insular environments; hence, density-reduction campaigns are often applied to alleviate the predation mortality they add to native fauna. Density-reduction and eradication efforts are costly procedures with important outcomes for native fauna recovery, so they require adequate planning to be successful. These plans need to include empirical density-reduction models that can guide yearly culling quotas, and resource roll-out for the duration of the culling period. This ensures densities are reduced over the long term and that no resources are wasted. We constructed a stochastic population model with cost estimates to test the relative effectiveness and cost-efficiency of two main culling scenarios for a 10-year eradication campaign of cats on Kangaroo Island, Australia: (1) constant proportional annual cull (one-phase), and (2) high initial culling followed by a constant proportional maintenance cull (two-phase). A one-phase cull of at least 0.35 of the annual population size would reduce the final population to 0.1 of its original size, while a two-phase cull with an initial cull of minimum 0.6 and minimum 0.5 maintenance cull would reduce the final population to 0.01 of its initial size by 2030. Cost estimates varied widely depending on the methods applied (shooting, trapping, aerial poison baits, *Felixer*™ poison-delivery system), but using baiting, trapping and *Felixers* with additional shooting to meet culling quotas was the most cost-effective combination (minimum cost: AU$19.56 million; range: AU$16.87 million–AU$20.69 million). Our model provides an adaptable and general assessment tool for cat reductions in Australia and potentially elsewhere, and provides relative culling costs for the Kangaroo Island programme specifically.

## Introduction

Since its domestication approximately 10,000 years ago, the common house cat *Felis silvestris catus* has spread throughout the globe and become established in most habitat types (including on most islands) (Fitzgerald et al. 1991; Medina et al. 2011; Woinarski et al. 2015), due to both accidental and deliberate human facilitation (Driscoll et al. 2007). Because they are generalist predators, feral cats are today one of the most destructive invasive mammal predators worldwide (Lowe et al. 2000; Doherty et al. 2016a), contributing to many of the predation-induced terrestrial (mainly island) extinctions recorded globally (e.g., > 63 species, including 26% of bird, mammal and reptile extinctions) (Doherty et al. 2016b).

The most effective method for removing the predation mortality on native species caused by feral cats is eradication wherever possible (Andersen et al. 2004; Schmidt et al. 2009), particularly in insular environments (Bester et al. 2002; Nogales et al. 2004; Doherty et al. 2016a). Alternative non-lethal approaches (such as trap-neuter-release) also exist (Gibson et al. 2002; Wallace & Levy 2006; Longcore et al. 2009; Miller et al. 2014), and while such an approach might appeal to members of the public that do not agree with lethal control (Andersen et al. 2004), the high expense of broad-scale implementation, coupled with its relatively low effectiveness compared to lethal methods (Longcore et al. 2009; Campbell et al. 2011) mean it is not widely used for cat management in Australia. Despite this, the trap-neuter-release management option is commonly considered in density-control programs, or proposed by communities (Deak et al. 2019).

Lethal control methods include poison baiting, trapping, and hunting (Campbell et al. 2011; DIISE 2018). *Eradicat*^®^ and *Curiosity*^®^ are poison baits developed specifically to target cats in Australia (Algar et al. 2011). *Curiosity*^®^ contains a robust, acid-soluble polymer pellet of para-aminopropiophenone poison (as opposed to 1080 poison, commonly used in dog and fox baits) (Department of Primary Industries and Regions 2020; Sharp & Quinn 2020), and is the only bait approved for feral cat control in South Australia (Department of Primary Industries and Regions 2020). Additionally, new technology is emerging in the field of feral cat baiting — particularly in terms of bait delivery — such as the *Felixer*™. The *Felixer*™ is an automated toxin-delivery system that uses rangefinder sensors to distinguish target cats from non-target species and sprays targets with a measured dose of toxic gel (thylation.com). Two types of trapping are often used simultaneously and in combination with baiting: cages and padded leg-hold traps. Animals are live-caught in traps and humanely dispatched, primarily with a 0.22-calibre rifle (Algar et al. 2020). Hunting is a term used for locating and shooting feral cats either during the day or at night (with the aid of a spotlight) from a slow-moving vehicle or on foot with a 0.22-calibre rifle (Nogales et al. 2004; Sharp 2018). Because shooting is the tool used for control, we refer to this technique as ‘shooting’ hereafter.

Most density-reduction campaigns based on direct killing have been typically implemented *ad hoc* because of the ongoing predation by cats on native prey species, and the requirement to achieve outcomes quickly (Bester et al. 2002; Denny & Dickman 2010). As such, available funds or resources can be used up quickly without the benefit of long-term planning based on the projections of empirical density-reduction models (Denny & Dickman 2010), thus threatening the success of a program. As a result, inappropriate methods and poorly timed roll-out have been attributed to most island eradication failures (Campbell et al. 2011). Custom-designed culling models that plan the most efficient and cost-effective application of resources are therefore ideal precursors to any eradication program (Smith et al. 2005; McMahon et al. 2010).

Culling models can be effective in this manner because of their ability to consider real-time population dynamics and resource availability to recommend feasible density-reduction plans (McMahon et al. 2010). Multiple types of culling model exist (e.g., spatially explicit, aspatial, density-driven, area-dependent) depending on the choice of scenario to reduce the population, such as a single-phase, constant proportional culling (McCarthy et al. 2013), or a two-phase cull with a high initial proportional culling rate followed by a constant proportional maintenance thereafter (i.e., a two-phase reduction model) (Campbell et al. 2011). Such models are instrumental in guiding successful eradication by providing targets and parameters that lead to efficient population reduction of the target species (Smith et al. 2005). Such two-phase eradication strategies (high initial cull followed by a consistent maintenance cull to ensure continued population decline) often still require a final ‘clean up’ stage where different (and usually more expensive) strategies are needed to eradicate the last surviving individuals that are difficult to detect (Bester et al. 2002; Nogales et al. 2004), and a ‘monitoring for success’ phase to ensure all animals have been removed (Algar et al. 2020). High initial culls followed by maintenance culling capitalise on the notion that when densities are high, culling is more efficient, while the maintenance culling continues to reduce the population as densities decline (Nogales et al. 2004; Denny & Dickman 2010).

Cat removal with the goal to eradicate is currently underway on part of Kangaroo Island, Australia’s third-largest island. The initial planning stages of the eradication began in 2016, with a proposed completion date of 2030. Kangaroo Island is a good candidate for eradication because of the island’s relatively intact native biodiversity compared to the mainland, and high local endemism (Taggart et al. 2019), as well as local community support for removing feral cats (Berris et al. 2019). Kangaroo Island’s feral cat eradication is part of the *Threat Abatement Plan*, that aims to “… prevent feral cats from occupying new areas in Australia and eradicate feral cats from high-conservation-value islands” (Environment 2015).

The program directors plan to use four main techniques for cat eradication on Kangaroo Island: baiting, trapping, shooting, and *Felixer*™ units. The social licence to apply lethal population reduction via shooting, baiting, or trapping is largely a function of the public’s perception of the proposed methods (Deak et al. 2019), and relies on the co-operation of land owners. This applies to Kangaroo Island given it has more than 4200 permanent residents spread across most of the island. Perception surveys done between 1993 and 2018 showed that > 90% of the Kangaroo Island community supported domestic and feral cat management (Berris et al. 2019).

Our aim was to design an ideal set of culling conditions that will most efficiently reduce feral cat densities on Kangaroo Island. More specifically, we (1) constructed stochastic variants of both culling and fertility-reduction (trap-neuter-release) models under different application scenarios that can be applied to guide cat eradication on Kangaroo Island, (2) estimate the relative costs of employing different combinations of the methods available, and (3) use the culling model to identify a regime that will most effectively reduce the feral cat population by the 2030 deadline. Specifically, we tested the efficacy (proportion of the population reduced, and over what time) of two culling scenarios: (*i*) constant proportional annual culling (one-phase), (*ii*) high initial culling followed by a constant maintenance cull (two-phase). We hypothesise that the two-phase culling model will reach the target population density by 2030 more efficiently than the one-phase culling model because initial effort tends to be the cheapest and most effective means of achieving high rates of reduction (Bester et al. 2002; Nogales et al. 2004; Robertson 2008; Denny & Dickman 2010). In any case, maintenance culling is required thereafter to prevent the population from recovering.

## Methods

### Study site

Located approximately 12 km south of the Fleurieu Peninsula (South Australia) at its nearest point, Kangaroo Island is Australia’s third largest island (155 km long and 55 km wide), covering ~ 440000 ha (Masters et al. 2004; Higgins-Desbiolles 2011) (Fig. 1). The island has retained around 53% of its native vegetation, with 35% of the remaining land cover devoted to dryland agriculture (Willoughby et al. 2018). The island is absent of invasive red foxes (*Vulpes vulpes*) and European rabbits (*Oryctolagus cuniculus*). As a consequence of the absence of rabbits, resident cats feed on a wider range of native species than elsewhere on mainland Australia (Bonnaud et al. 2011).

**Figure 1.**
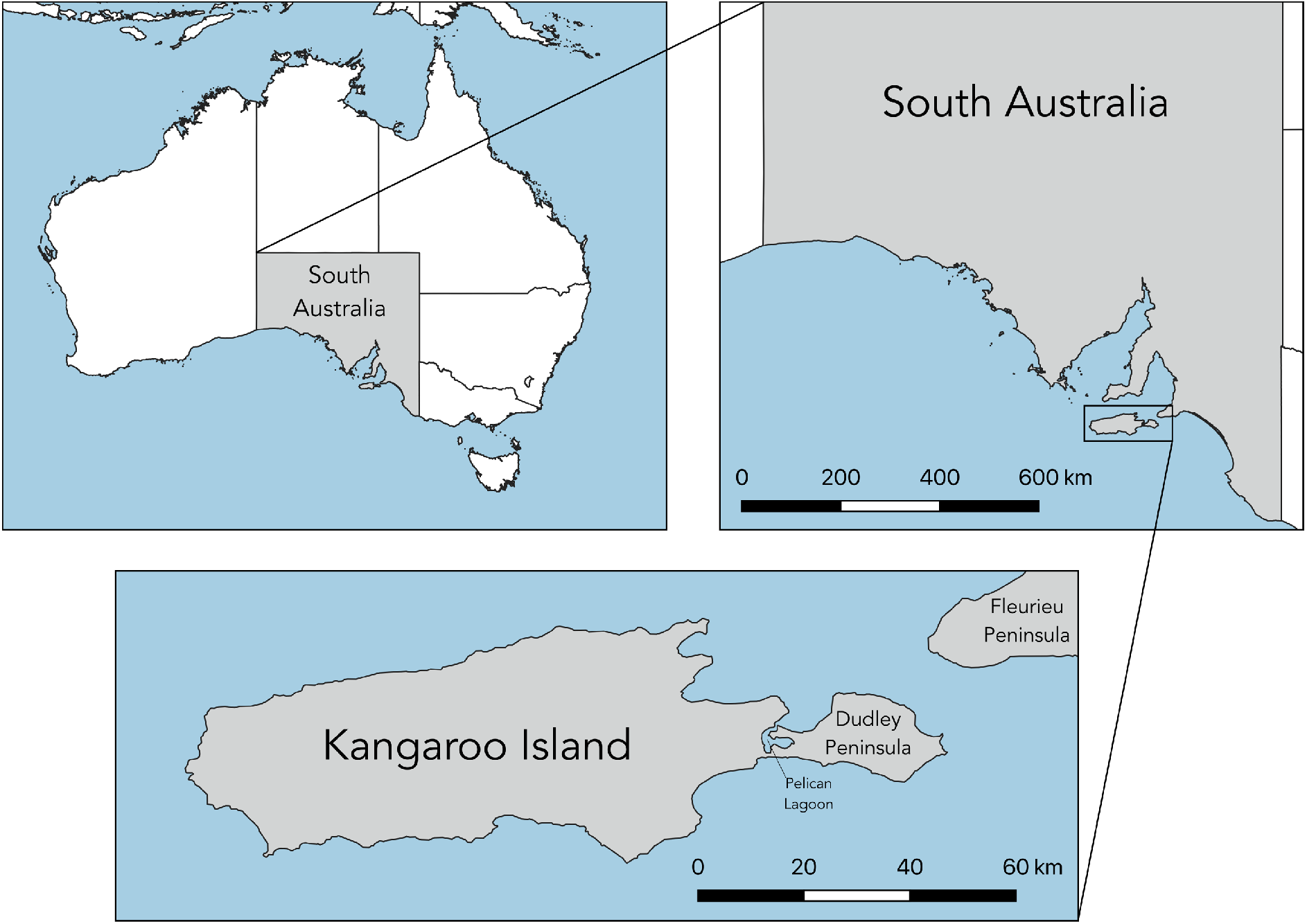
Map of Kangaroo island relative to the Australian mainland. The shortest distance from the mainland (southern tip of Fleurieu Peninsula) to Kangaroo Island is approximately 14 km.

Feral cat densities on Kangaroo Island are thought to range from 0.06 to 3.27 cat km^−2^, with an average density of 0.37 cat km^−2^, giving an estimated population size of 1629 (s.e. ± 661) individuals (Hohnen et al. 2020a; Hohnen et al. 2020b). Taggart et al. (2019) estimated that relative feral cat densities in eastern Kangaroo Island were ~ 10 times higher than on the adjacent mainland (Kangaroo Island relative abundance = 14.6 cats camera-trap-site^−1^; mainland = 1.39 cats site^−1^; 11 sites on both the Island and mainland).

### Model

We constructed a Leslie matrix to represent age-specific fertility and survival (Caswell 2001) for the cat population on Kangaroo Island. We obtained cat fertility and survival estimates from six studies of domestic, stray and feral cat population across the USA and Australia (Budke & Slater 2009), and summarised the population dynamics from a study in Western Australia done in the preliminary stages of cat eradication (Short & Turner 2005). We calculated mean and standard deviations of the age-specific demographic rates (i.e., survival, fertility) necessary for stochastic representations of the model (see below). We only used these fertility and survival estimates for females, assuming a 1:1 sex ratio (Bloomer & Bester 1991; Budke & Slater 2009).

According to demographic rates published in the peer-reviewed literature, the maximum age for feral cats ranges from 3 (Budke & Slater 2009) to 9 years (Van Aarde 1983). We set the maximum age to the median maximum age in the literature: 6 years; this was supported by feedback from wildlife managers on Kangaroo Island. Cats become sexually mature between 6 and 12 months of age (Jemmett & Evans 1977; Povey 1978; Jones & Coman 1982; Bukowski & Aiello 2011). To account for pre-yearling reproductive output, we reduced the fertility in the initial year by one-third to represent the approximate proportion of juveniles breeding. For all resulting predictions of changing population size, we assumed that survival was the same for males and females given no evidence to the contrary. We present all parameters and their ranges in Table 1.

**Table 1.** Mean parameter values and their standard deviations (SD) used in the stochastic model.

| **parameter** | **mean** | **SD** |
| --- | --- | --- |
| <u>*fertility (daughters)*</u> |  |  |
| pre-breeding* juvenile ($f_1$) | 0.000 | — |
| sub-adult ($f_2$) | 0.745 | 0.307 |
| adult ($f_3$–$f_7$) | 2.520 | 0.450 |
| <u>*survival*</u> |  |  |
| pre-breeding juvenile ($s_1$) | 0.460 | 0.115 |
| sub-adult ($s_2$) | 0.460 | 0.115 |
| adult ($s_3$–$s_6$) | 0.700 | 0.058 |
| senescent ($s_7$) | 0.550 | 0.058 |

\*pre-breeding juvenile: < 10 months old; not sexually mature

We stochastically resampled at each time step in the deterministic matrix **A** in all subsequent projections based on the standard deviation estimated from minimum and maximum fertility and survival values (Budke & Slater 2009), which incorporates both measurement error and inter-annual variability (process error). We assumed a Gaussian distribution around the mean of fertility and the *β* distribution for survival probability, using the standard deviations for resampling of each (Table 1).

The deterministic matrix **A** is:

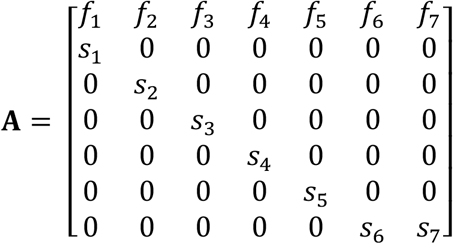

where *f_x_* = age (*x*) -specific fertility and *s_x_* = age-specific survival (note: *f*_1_ represents 0 years, because individuals are < 1 year old in their first year). See also Table 1 for parameter values.

### Untreated (control) population

To simulate how incrementing intensities of reduction alter the projected population size, we first simulated a population not exposed to any culling to represent a ‘control’ population. We calculated the population’s stable age distribution from the base matrix **A** (Caswell 2001), and then multiplied this stable age structure by a starting population size of 1629 (Hohnen et al. 2020b). We then expressed all subsequent projections as a proportion of this founding population size to avoid the uncertainty in initial population size estimates. Kangaroo Island is insular, so there are fewer opportunities for migration into the population compared to the mainland, and local residents are largely cooperative with the regulation of domestic cats to assist in eradication (> 90% community support; Berris et al. 2019). However, we did account for some ‘leakage’ into the population (domestic release or ferry stow-away) by re-running the top-performing culling scenario and adding an incrementing number of ‘leaked’ individuals (between 10 and 1000 cats) into the population annually to test how these additions would affect final population sizes post-culling (see Supporting Information, Appendix A, Fig. S1).

We included a logistic compensatory density-feedback function by reducing survival when the population exceeded double the size of the current population (see below) of the form:

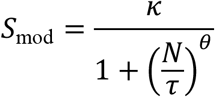

where *S*_mod_ is the proportion of realised survival (survival modifier) as a function of the population’s proximity to carrying capacity (twice the founding population size, *K* = 3258; see below), *N* is the population size, and *κ, τ* and *θ* are constants: *κ* = 1.001, *τ* = 5459.994, and *θ* = 1.690 (see Supporting Information, Appendix B, Fig. S2). We thus assumed that survival probability would decline as the population approached carrying capacity (double the size of the current population). The feedback mechanism means that as the population approaches carry capacity, survival across all ages is reduced by *S*_mod_ according to this relationship. This function acts to drive total population size away from carry capacity. We set carry capacity to twice the initial population because landscape managers currently consider the population to be below carry capacity with respect to available food resources (Jones & Coman 1982; Read & Bowen 2001). While the carrying capacity is somewhat arbitrary, it does realistically allow the population to increase if no additional mortality sources are imposed. Most research on feral cat population control does not consider the habitat’s carrying capacity (Andersen et al. 2004); however, feral cats seem to maintain consistent fertility regardless of population density, although average survival tends to decrease as the population approaches carrying capacity (Courchamp & Sugihara 1999; Nutter 2006). We therefore did not adjust fertility relative to population size.

### Reduction scenarios

#### 1. Trap-neuter-release

To compare the efficacy of our modelled density-control and -reduction scenarios with fertility-reduction methods, we constructed a model that simulated a trap-neuter-release implementation. Although not widely used, trap-neuter-release is often suggested by a certain element of the public as a more ethical alternative to lethal control. We included this scenario here to compare its efficacy directly to the culling scenarios described below. In this model, no animals are removed from the population, but fertility is reduced to simulate sterilisation. We ran this model using the same methods for an unculled population (and over the same interval), but we reduced fertility for each iteration across a range of values (1–99%, at 1% intervals; e.g., fertility reduced by 50% in one scenario 51% in the next, and so forth). This represents the percentage of the population that is neutered (neutered individual fertility = 0), giving a realised population fertility of between 99% and 1% of non-intervention values (depending on the pre-determined fertility-reduction target of the scenario). Each year, new individuals are neutered to maintain the predetermined population fertility.

#### 2. Culling model

We built two culling models: (*i*) constant proportional annual culling (one-phase), and (*ii*) a high initial proportional cull in the first two years, followed by a constant proportional maintenance cull (two-phase). Here we consider only these phases of a strategy where the last step likely requires a ‘clean-up’ — the latter is difficult to consider in a model because the focus shifts to individuals. Additionally, we did not consider ‘monitoring for success’ as this stage does not involve culling *per se*. We instead defined a threshold at the end of the model, where moving to the ‘clean-up’ stage is deemed feasible. In each model, we removed individuals from the population vector proportional to the total culling invoked in that time step and the stable age distribution. We ran each model for 10,000 iterations (randomly sampling 10,000 times from the stochastic survival and fertility vectors) to calculate the mean and 95% confidence bounds for minimum proportional population size. We set the projection interval to 10 years to represent the management strategy for eradication by 2030 (2020–2030).

For the one-phase scenario, we simulated constant proportional annual culling (*c*) (i.e., we reduced the population each year by the same proportion for the duration of the projection interval) from *c* = 0.20 to 0.90, at intervals of 0.05. For the two-phase scenario, we applied high initial culling only in the first two years of the eradication project (*c* = 0.50–0.99), with maintenance culling applied to all years thereafter (*c* = 0.01–0.50) until the end of the projection interval. For all iterations of both models, we recorded the minimum projected proportional population size (p*N*) for each value of *c*, at an incrementing proportional culling of 0.01.

### Cost

Based on previous information regarding the reduction in capture efficiency as population density declines (Bloomer & Bester 1992; Nogales et al. 2004; Parkes et al. 2014), we assumed an eradication technique’s efficiency (*f*, ranging from 0 to 1) follows a Type III functional response (i.e., sigmoidal; Nunney 1980, Denno and Lewis 2009) relative to proportional population size:

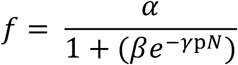

where *f* is the relative efficiency of the culling technique, *α*, *β* and *γ* are constants: *α* = 1.01, *β* = 85.61, and *γ* = 8.86, and p*N* = proportional population size (see Supporting Information, Appendix C, Fig. S3). We assumed the same efficiency reduction across trapping, shooting, baiting, and *Felixers*™ as a function of population size, such that the smaller the remaining population of cats, the less efficient each method was relative to the start of the eradication campaign. We then applied this reduction to the culling model with pre-set costs for each technique (see below), to estimate the total cost of eradication.

We sourced trapping and shooting cost data from Holmes et al. (2015), with additional trapping costs from trapping supplies (traps.com.au) and *Felixer*™ data from Moseby et al. (2020). We sourced aerial baiting data from Johnston et al. (2014) and Algar et al. (2020), as well via direct correspondence with the Australian federal Department of Agriculture, Water and the Environment (Julie Quinn, Canberra, Australian Capital Territory, pers. comm.) and Wrightsair (wrightsair.com.au; Ellodie Penprase, William Creek, South Australia, pers. comm.).

We summarised the catch rates and costs for each technique: (*i*) *Felixer*™ — each unit costs AU$13,000. Based on efficacy trials at Arid Recovery, 20 *Felixer*™ units deployed over 2,600 ha were successful in killing 31 cats over 41 days, which translates to an annual kill rate of 5.749 (cats killed unit^−1^ year^−1^). (*ii*) Traps — each trap costs between AU$157 and AU$297 (sampled uniformly). Based on trials on Dudley Peninsula, 40 traps deployed over approximately 12,000 ha caught 21 cats in 148 days, which translates to an annual trap rate of 0.198 cats trap^−1^ year^−1^. (*iii*) Shooting — from Holmes et al. (2015), we estimated a kill rate person-hour^−1^ based on 1044 kills (872 direct + 172 from wounds) over 14,725 person-hours (= 0.071 cats killed person^−1^ hour^−1^). Ammunition and labour costs equate to AU$25.92 hour^−1^. (*iv)* Baiting — each *Curiosity*^®^ bait costs $2.27 unit^−1^, with a one-off AU$250 administration fee order^−1^ (treidlia.com.au; Arsalan Shah, Tréidlia Biovet Pty. Ltd., Seven Hills, New South Wales, pers. comm.). We received fixed-wing charter costs directly from Wrightsair that quoted AU$750 hour^−1^ when actively baiting and AU$600 hour^−1^ for chartering aircraft from their base in William Creek, South Australia. From Johnston et al. (2014) based on 15 collared cats, an average density 0.701 cats km^−2^ (approximate total area: 15 × 0.701 = 10.515 km^2^), with 50 baits km^−2^ (526 baits), killed 14 cats (= 0.026 cats bait^−1^ or 37.55 baits cat-killed^−1^).

To estimate total costs, we first assumed that the density of traps applied on Dudley Peninsula (Dudley Peninsula = 37,500 ha) could be extrapolated to the much larger area of the entire island (440,500 ha). Based on these densities, we calculated the total number of traps required for the entire island, and then tabulated the number of cats killed by this method for the incrementing proportional cull. We then combined this with baiting for the initial phase of culling.

If the total number of cats killed by these methods fell short of the proportional cull target in any given year and iteration, we applied three different scenarios where we varied the method used to achieve the proportional target beyond the initial roll-out of units and traps. The three different approaches to meet the shortfalls were: (*i*) *Felixers*™, (*ii*) increasing the number of traps only, or (*iii*) meeting the shortfall entirely with follow-up shooting. In each shortfall scenario, we tabulated the total costs across the projection interval and expressed these as a function of the increments in proportional culling.

Of course, this approach assumes a simultaneous roll-out of all *Felixer*™ units and traps across the entire island, when a more efficient approach might instead be to purchase a smaller number of units/traps and deploy them in a spatially sequential roll-out (i.e., a moving ‘wave’ of units applied to specific regions of the island in sequence as localised eradication is achieved). We therefore also ran a modified scenario to reflect this type of spatial pattern of application by arbitrarily assuming a smaller number of units/traps across the entire landscape. Reducing the purchase cost per unit/trap by the same arbitrary value is therefore functionally equivalent to a spatially sequential roll-out of this smaller sample of units/traps. For this example scenario, we therefore reduced the purchase cost of both *Felixers*™ and traps by two-thirds unit^−1^ (see Supporting Information, Appendix D, Fig. S4).

## Results

### Untreated population

An untreated (no-cull) ‘control’ population is expected to increase to a median of 1.9 times the founding population (i.e., to 3118 individuals when starting with 1629) by 2030 (95% confidence limits: 0.919–3.324 times) (Fig. 2a). The instantaneous rate of change (*r*) from the deterministic matrix for the Kangaroo Island population is 0.222. The deterministic (mean) matrix gave a generation length of 3.207 years. The population is projected to approach carrying capacity (set arbitrarily at twice the current population size) and begin to plateau by 2028 (Fig. 2a), at which time the population’s median *r* from the stochastic projections is 0.009. By 2030, the population’s median *r* from the stochastic projection is 0.005.

**Figure 2.**
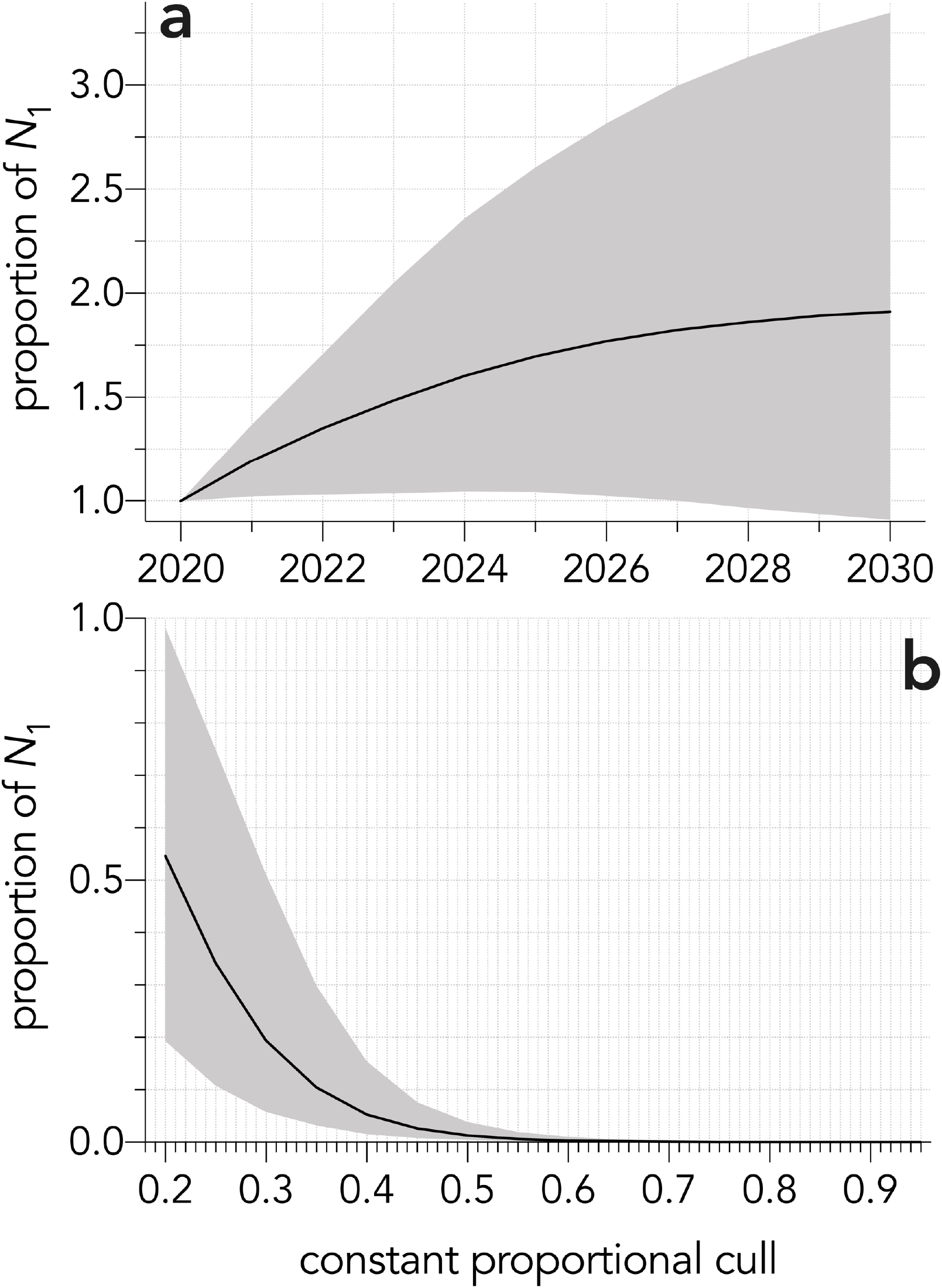
(a) Average proportion of the initial cat population (*N*_1_) on Kangaroo Island projected from 2020–2030 for the unculled scenario. Black line indicates the median value from 10,000 iterations, along with 95% confidence intervals (grey-shaded area). (b) Minimum proportion of the Kangaroo Island feral cat population remaining after a constant proportional annual cull ranging from 0.2 to 0.9. Solid black line represents median minimum proportion of the initial population (*N*_1_) after 10,000 iterations with 95% confidence intervals indicated as grey-shaded area.

### Trap-neuter-release

The trap-neuter-release scenario would reduce the population to < 0.1 (95% confidence limits: 0.036–0.221) of its original size by 2030 when the population’s overall fertility is reduced by 26% (Fig. 3). A 55% reduction in fertility would drive the population to < 0.01 (0.021–0.003) of its original size by 2030.

**Figure 3.**
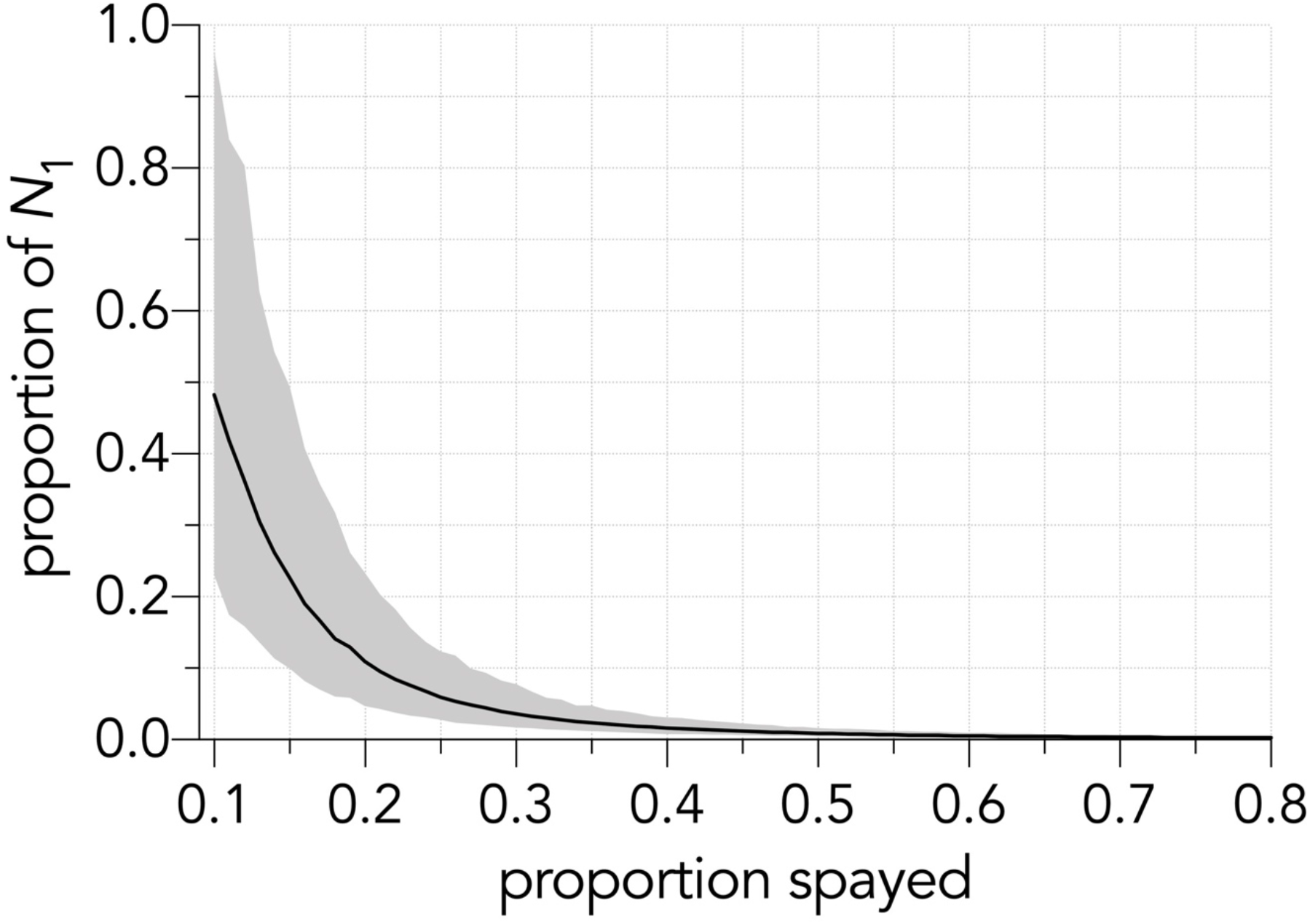
Estimated median minimum proportion of founding population remaining (founding population *N*_1_), with decreasing fertility (fertility reduced 50–99%). Solid black line represents median minimum proportion of the initial population (*N*_1_) remaining after fertility reduction scenarios, with 95% confidence intervals indicated as grey-shaded area.

### Culling

Culling timeline 2020–2030: For the one-phase scenario, a minimum annual proportional cull of 0.35 would reduce the population to 0.10 of its initial size. A minimum annual cull of 0.5 would reduce the population to 0.01 of its initial size (Fig. 2b). A two-phase cull with a minimum initial cull of 0.55 and a minimum maintenance cull of 0.3 would reduce the population to 0.10 of its initial size. To reduce the population to 0.01 of its initial size requires a minimum initial cull of 0.60 followed by a minimum maintenance cull of 0.5 (Fig. 4) Stopping the program after the initial culling during the first two years (i.e., without any maintenance culling), the population would recover to its initial size in 15 years (range: 11–21 years; Fig. S4), whereas stopping the maintenance cull in the 9^th^ year (i.e., a year before termination of the program) would result in population recovery to initial size in 42 years (range: 35–50 years) (Fig. S4). ‘Leakage’ from stray cats had little overall effect on the effectiveness of the total cull (Supporting Information, Appendix A, Fig. S1).

**Figure 4.**
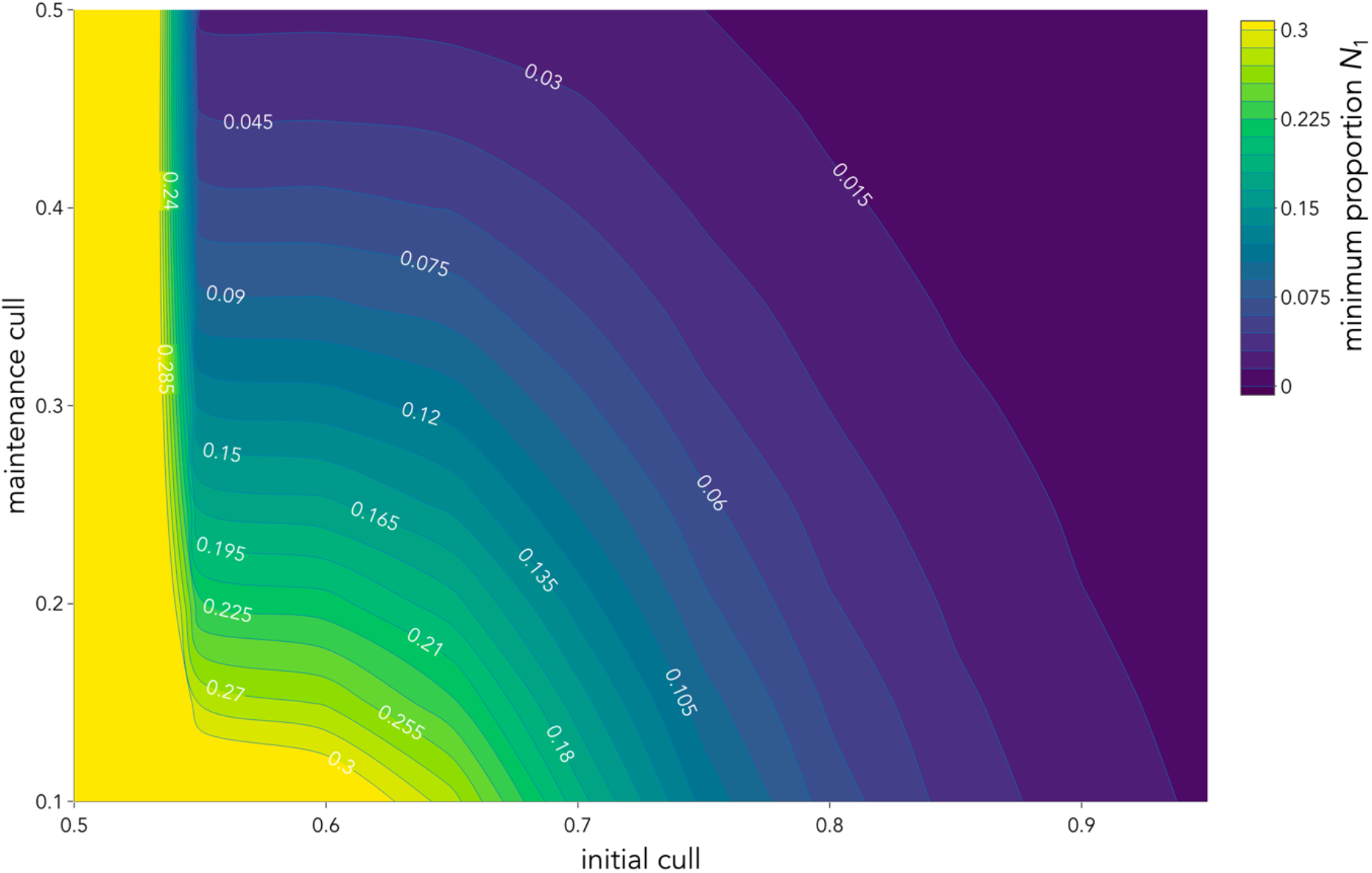
Estimated median minimum proportion of the final population remaining (relative to start population *N*_1_) for combinations of initial proportional (i.e., *initial cull*: 0.5–0.9) and maintenance proportional (i.e., *maintenance cull*: 0.1–0.5) culling. Proportion of population remaining after culling scenarios represented by colour bar ranging from lowest (purple) to highest (yellow) remaining proportional population.

### Cost

To reduce the entire Kangaroo Island population to a 0.10 of its original size (0.1*N*_1_) using a two-phase cull (minimum 0.55 initial, 0.3 maintenance), a minimum of AU$19.56 million (AU$16.87 million–AU$20.69 million) (Fig. 5c), would be required if shooting was used to make up the yearly shortfall. In contrast, making up the shortfall with additional traps would increase the average costs by 88.75% to AU$36.92 million (AU$27.07 million–AU$47.27 million) for the same target (Fig. 5b). Finally, making up the shortfall with additional *Felixer*™ units would increase the average cost relative to the shooting-shortfall scenario by 226.6% to AU$63.89 million (AU$47.56 million–AU$70.17 million) (Fig. 5a). Changing the target population size to 0.01 (minimum 0.60 initial, 0.5 maintenance) of the initial (0.01*N*_1_), the total minimum costs would increase to AU$24.38 (AU$21.96–AU$27.29 million) if the shortfall was made with shooting (24.64% more than the 0.1*N*_1_ shooting-shortfall scenario), AU$52.60 million (AU$38.69 million–AU$70.26 million) if the shortfall was made with traps (115.7% more than the 0.01*N*_1_ shooting-shortfall scenario), or AU$93.65 million (AU$78.52 million–AU$1.11 billion) if the shortfall was made with *Felixers*™ (284.12% more than the 0.01*N*_1_ shooting-shortfall scenario) (Fig. 5).

**Figure 5.**
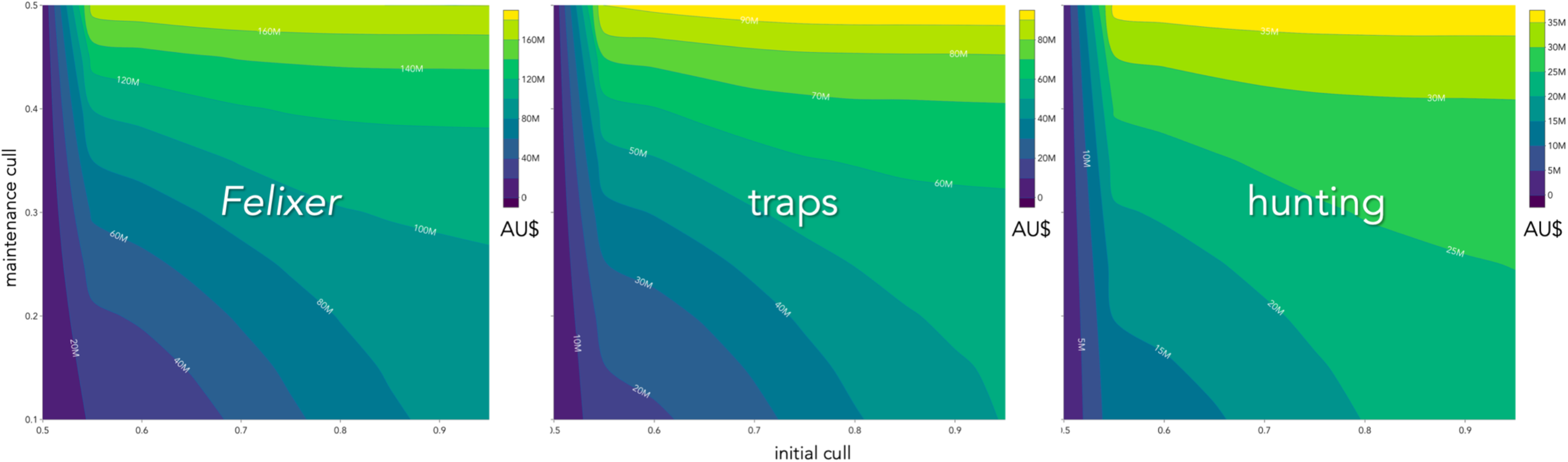
Estimated median total costs of feral cat eradication on Kangaroo Island for combinations of initial proportional (i.e., *initial cull*) and maintenance proportional (i.e., *maintenance cull*) culling, where the shortfall in the number of cats killed from *Felixer*™ units and traps is provided by (a) additional *Felixer*™ units, (b) traps, or (c). Cost of eradication (in AU$, adjusted for 2020) indicated by colour bar ranging from lowest (purple) to highest (yellow) costs. Contours and white values indicate cost in $AU millions. Note different *z*-axis (contour) scales in a, b, and c.

## Discussion

In each cull scenario we considered, a successful reduction of the feral cat population on Kangaroo Island to below 0.01 of its initial size by 2030 is achievable, but the minimum costs involved according to the different scenarios we ran could range from AU$24.38 million (AU$55 ha^−1^, with shooting; Fig. 5c) to AU$93.65 million (AU$213 ha^−1^, with *Felixers*™; Fig. 5a), depending on the method used and the inherent uncertainty in the parameters we estimated.

The realism of our modelled total cost estimates depends on the form of the (as-yet unmeasured) functional response, and the assumed per-unit efficacy of the eradication tools to meet the annual shortfall for the predetermined cull proportion. Reported costs for feral cat island eradications globally have a large range (AU$6 ha^−1^ – AU$314 ha^−1^; adjusted to 2021 AU$; Campbell et al. 2011). Our lower cost estimate for culling only (AU$55 ha^−1^) is 800% greater than the cost of complete eradication (including clean-up and monitoring for success) on Faure Island, Western Australia (AU$6 ha^−1^) (Algar et al. 2010). However, Faure Island covers 5800 ha — and is therefore only 1.3% the size of Kangaroo Island (440000 ha). Dirk Hartog Island (62000 ha) is larger, at 14.1% the size of Kangaroo Island, and is currently the largest successful island eradication globally (Algar et al. 2020). Eradication there cost approximately AU$90 ha^−1^, although that included construction of a barrier fence, clean-up, and monitoring for success (Algar et al. 2020). Our cost estimates for Kangaroo Island are 377% cheaper than the cost to remove cats from Macquarie Island (AU$258 ha^−1^), likely due to the latter’s remoteness (Robinson & Copson 2014). Cost estimates are notably underreported in the literature; Campbell et al. (2011) found < 10% of successful island eradications reported costs. Further, reported costs are often whole costs, and provide little detail into money spent per stage (culling, clean-up, monitoring for success), making direct comparisons difficult.

Nonetheless, our outputs do suggest that high initial culls (> 0.55, 0.6) followed by moderate maintenance culls (0.3–0.5) would be sufficient to reduce the population to 0.01–0.10 of its original size (Fig. 4), and that shooting is the most cost-effective way to meet these targets (especially if other methods are rolled out simultaneously).

That shooting is cheaper than other methods is unsurprising given that no hardware other than rifles and ammunition is needed to be purchased outright, in contrast to the higher overheads associated with traps or *Felixer*™ units (Holmes et al. 2015; Hodgens 2019). However, it is not reasonable to assume that the *Felixer*™ will be used in the same widespread capacity as trapping and shooting. The *Felixer*™ is more likely to be used sporadically, or to target areas that are not appropriate for trapping or shooting (e.g., roadsides, thick bushland, areas definitely known to be frequented by cats) (Moseby et al. 2020). Therefore, our cost estimates using *Felixers* to make up the shortfall are for comparison rather than being recommendations *per se*.

Indeed, shooting requires many people working full time, whereas the other techniques are more passive (yet the latter also require set up, monitoring, maintenance, displacement, and removal by staff). However, approximately 500 person hours are required to equate to the current cost of a single *Felixer*™ unit (shooting ~ AU$26 per person hour^−1^ vs. AU$13,000 *Felixer*™ unit), although assuming a spatial roll-out of fewer units is functionally identical to a similar reduction in per-unit cost. Additionally, shooting is considered more humane due to the minimised contact with the animal and the instant death with a correctly executed headshot (Sharp & Saunders 2011), but access to private land and potential conflict with private landholders could complicate shooting because of the social licence needed for lethal control. Of course, complete eradication would necessarily entail additional costs as the final individuals were identified, hunted, and destroyed (Bester et al. 2002; Nogales et al. 2004), and a monitoring stage to ensure all individuals are removed (Campbell et al. 2011; Algar et al. 2020). These final stages are important because even a few individuals remaining could conceivably seed a recovery that could achieve initial population size in several decades (Supporting Information, Appendix D).

Our results also identify that fertility-reduction using trap-neuter-release methods are comparatively ineffective for reducing pest densities (Longcore et al. 2009). Our model output suggests that the population would need to have a realised fertility of 74% for the entirety of the study period (2020–2030) to reduce it below 0.1 of the initial population. Whether fertility reduction is feasible or cost-effective is beyond the scope of our study, but it does demonstrate that fertility-reduction is a much less efficient method to eradicate cats than culling. That trap-neuter-release is less efficient than lethal control is not a new finding. Matrix modelling for a free-roaming cat colony found population reduction to be more feasible with euthanasia than sterilisation (Andersen et al. 2004). Further, efforts to remove urban feral cats in Hawaii found the trap-neuter-release method less cost-effective than lethal control, even when the former employed volunteers and the latter employed paid professionals (Lohr et al. 2013). Finally, Campbell et al. (2011) found no successful eradications on islands using the trap-neuter-release method. Additionally, sterilised individuals returned to the population could still continue to eat native fauna until they perished due to natural causes, so the risk cats pose to their prey is not diminished instantaneously, as it is with culling-based programs (Andersen et al. 2004).

We conclude that the most appropriate approach to reduce cat densities on Kangaroo Island is a two-stage method, with a high initial reduction of at least 0.55–0.7 and a maintenance cull of 0.3–0.65. Although a constant proportional annual cull can be effective, it is generally less efficient than a two-stage approach. This is because effort is spread equally among temporal windows in the constant proportional scenario, and therefore must ‘catch up’ relative to a large, initial cull given that more surviving individuals are still breeding in the former. As culling reduces density and drives the population closer to extinction, it becomes progressively more difficult and expensive to cull remaining individuals (Nogales et al. 2004; Parkes et al. 2014). This is because most culling methods are passive and rely on a ‘non-negligible probability’ of the target animal encountering *Felixers*™, baits, or traps (Moseby & Hill 2011; Fancourt et al. 2021). Although these techniques can be accompanied by visual, scent, or sound lures, the target animal still needs to be in range to be enticed by them. Thus, encounters at low densities become increasingly less likely (Veitch 2001; Campbell et al. 2011), and rising per-capita food abundance as the predator’s population dwindles can make baits or food lures less attractive (Parkes et al. 2014). Aerial baiting is most effective in the initial years of eradication because they can be widely distributed, including in areas that are inaccessible with vehicles or on foot (Nogales et al. 2004; Parkes et al. 2014).

Cats in particular are intelligent predators and can learn to avoid traps and baits. Thus, while these methods are generally considered effective for density reduction, it is most effective in the early stages of eradication programs (Nogales et al. 2004). Therefore, a two-stage approach allows for the implementation of widespread control that is effective at high densities, followed by a more targeted approach through consecutive maintenance as the population continues to decline.

The merits of the stochastic framework we developed imply that the model is transferable to other regions and even other species. Altering locally measured demographic rates, population sizes, control effectiveness, and reduction targets are feasible with this approach. For example, the Australian federal government has prioritised Christmas Island (Australian territory), Bruny Island (Tasmania), and French Island (Victoria) for eradication (Bannister 2017). All three islands have permanent human residents (French Island: 110; Bruny Island: 800; Christmas Island: 1840) and are considered large (greater than 1000 ha; Nogales et al. 2004). Our model is also applicable to mainland density control and eradications; however, it is only recommended for eradication in exclusion zones because cats can rapidly recolonise areas that have undergone density reduction (Moseby & Hill 2011; Palmas et al. 2020). Our model also has applications for other species, including European red foxes (Edwards et al. 2004) in Australia (particularly mainland exclusion zones), brush-tailed possums (*Trichosurus vulpecula*) and stoats (*Mustela erminea*) in New Zealand (Brown et al. 2015), and mainland application for species such as racoons (*Procyon lotor*) in central Europe (Beltrán-Beck et al. 2012).

For effective eradication to be achieved, culling programs must be based on empirical data and ideally, directed by models like ours. Our model should allow practitioners to make their culling programs more efficient, and to allocate the resources needed to achieve their targets efficiently and cost-effectively. As more site-specific data become available, we expect the model’s predictions to become ever-more realistic to identify the most plausible and cheapest pathways to eradication.

### Perspective on the Kangaroo Island cat-eradication program

Due to insufficient data for many model parameters and functions, we were obliged either to make (arguably defensible) assumptions or use data from other locations/studies (Budke and Slater 2009). Although we are confident that our results and scenarios are relevant, they will undoubtedly be improved by the refinement of locally measured parameters such as age-specific demographic rates, updated density estimates following the 2020 bushfires, cost data, efficiency relationships, strength of compensatory density feedback, and probability of leakage. A more detailed schedule for resource application including budget restrictions, timeline flexibility, currently available resources (to reduce initial costs via unit purchasing) and available staff would also assist in improving the realism of the predicted scenarios. Further, the *Felixer*™ is still in the initial phases of production and is not yet produced on a large commercial scale. The *Felixer*™ has many merits in regards to feral cat control because of its hazard reduction for baiting non-target species (Read et al. 2019); thus, it is likely to be increasingly applied in future management projects, especially as costs per unit decline.

## Data availability

The R code to create the model simulations is available at https://github.com/KathrynVenning/FeralCatEradication.

## Acknowledgments

The authors acknowledge the traditional owners of the land we work on, and pay our respects to Elders, past, present and emerging. We thank the Kangaroo Island section of the South Australia Department of Environment and Water for data access and insights.

The R code described in this manuscript is available at https://github.com/KathrynVenning/FeralCatEradication

