## Supplementary material for "Predicting targets and costs for feral-cat reduction on large islands using stochastic population models"

Authors (redacted)

The following supplementary material includes figures, methods, results and further contextualisation for information given in the main text.

### Table of contents

**Appendix A – Leakage into the population**

**Appendix B – Survival modifier**

**Appendix C – Type III functional response**

**Appendix D – Stopping culling prematurely**

**Supplementary References**

### Appendix A – Leakage into the population

#### Background

The opportunities for immigration of cats to this island population are low because of strict controls on cat ownership and importation. However, accidental or deliberate releases, or domestic cats breeding with feral animals, are still plausible. All domestic cats on Kangaroo Island must be de-sexed, registered and microchipped by law (Natural Resource Kangaroo Island 2015), which greatly reduces the possibility of a domestic-feral breeding. Nonetheless, we considered a hypothetical scenario of 'leakage' of domestic cats into the population on Kangaroo Island to explore this phenomenon's effects on eradication efficiency.

#### Approach

We used the optimal-culling scenario (two-phase; initial 0.6, maintenance 0.5) to identify how leakage would affect the population at the end of the culling scenario. To achieve this, we added an incrementing number of individuals (between 0 and 100, in intervals of 10) annually to the population vector, while simultaneously running the culling scenario. We ran each leakage scenario for 10,000 iterations (randomly sampling 10,000 times from the stochastic survival and fertility vectors) to calculate the mean and 95% confidence limits for minimum population size.

#### Results

If no cats are leaked into the Kangaroo Island population (i.e., baseline expectation), the final population at the completion of culling would be 14 cats (range: 5–47, 95% confidence intervals). This would increase by a factor of 1.9 (i.e., 27 cats; range: 13–63) if 10 cats were added annually (Fig. S1). If 50 cats were added annually, the final population would increase to 79 cats (53–130) (Fig. S1).

**Figure S1**. Final population ($N$) at completion of the ideal culling scenario (two-phase; initial 0.60, maintenance 0.2), adding 0–100 cats annually into the population. Solid black line represents mean population after stochastic resampling. Grey shaded area represents 95% confidence bounds for minimum population size.

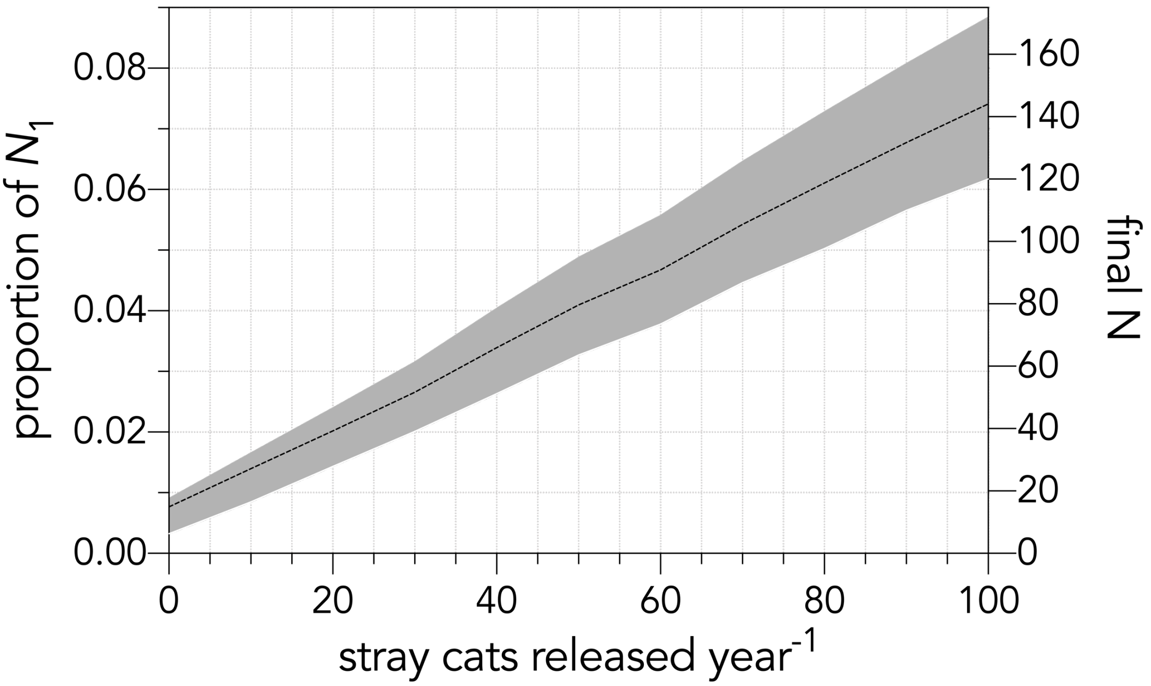


**Implications**

There are currently 244 registered domestic cats on Kangaroo Island, including a single breeder cat (presuming all cat owners follow the by-laws, this equates to only one domestic cat able to reproduce; Natural Resource Kangaroo Island 2015). It is therefore highly unlikely that 100 cats are leaked into the population annually. However, it is possible that a few cats (1–5) are accidently released annually. Likewise, it is possible some cat owners do not register their pets or accommodate 'stray' cats (provide food and/or water, but no direct ownership). Based on our simulation, leakage to this extent is unlikely to reduce overall eradication efficiency, particularly if owners abide by the by-laws, and with > 90% public approval for eradication (Natural Resource Kangaroo Island 2015) compliance with existing by-laws will be high. Regardless, even an annual leakage of 10 cats could increase the final population after the culling scenario to 1.9 times compared to the baseline scenario (Fig. S1). As such, wildlife practitioners and council members should continue community engagement and to enforce cat ownership by-laws.

### **Appendix B – Survival modifier** (compensatory density-feedback mechanism)

**Figure S2**. Reduction in probability of survival relative to population size (*N*), with carry capacity *K* set to twice the size of the founding population (3258). $\kappa$, $\tau$ and $\theta$ are constants ($\kappa = 1.001$, $\tau = 5459.994$ and $\theta = 1.690$).

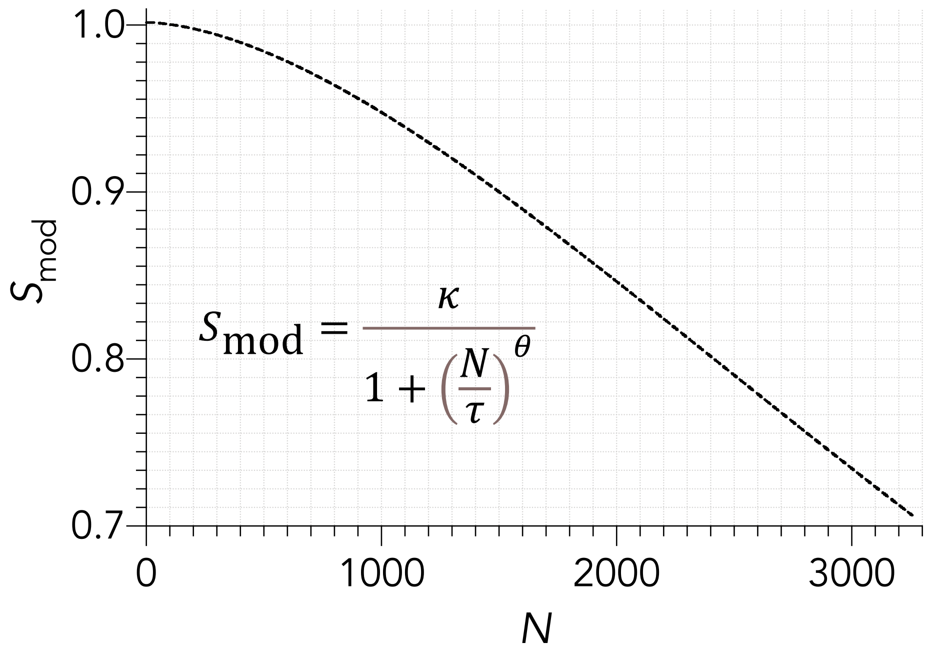

### Appendix C – Type III functional response

To force the model to adjust the efficacy of the different culling methods according to the number of individuals in the population, we constructed an arbitrary Type III functional response of the form:

$$f = \frac{\alpha}{1 + (\beta e^{-\gamma \mathrm{p}N})}$$

where $f$ is the relative efficiency of the culling technique, $\alpha$, $\beta$ and $\gamma$ are constants: $\alpha = 1.01$, $\beta = 85.61$, and $\gamma = 8.86$, and p$N$ = proportional population size (Fig. S3). There is no empirical basis for these particular parameter values; instead, we wanted to demonstrate a generic sigmoidal function to alter the efficiency of each culling method.

**Figure S3**. Reduction in the efficiency of a culling method ($f$) as the proportion of the remaining population size declines (p$N$). Here, $\alpha$, $\beta$ and $\gamma$ are constants ($\alpha = 1.01$, $\beta = 85.61$ and $\gamma = 8.86$).

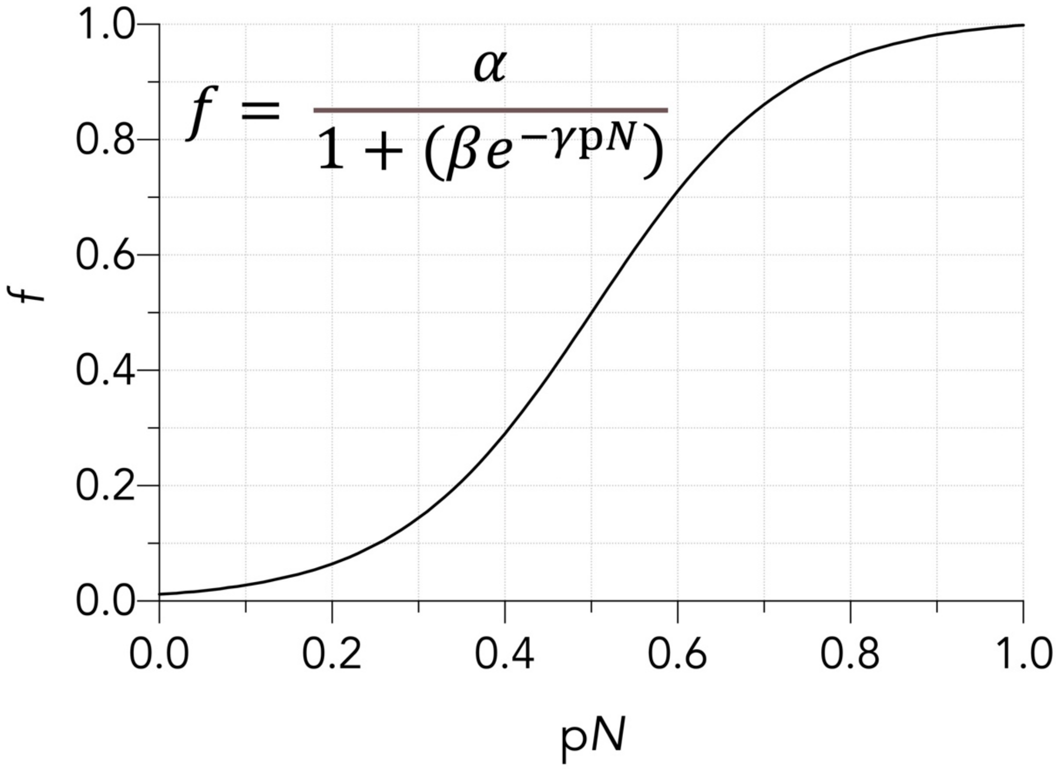

### Appendix D – Stopping culling prematurely

#### Background

Following eradications to completion are essential to achieve density-reduction or eradication goals, but due to funding limitations or unforeseen circumstances, such programs are sometimes cancelled or stopped prematurely (Keitt et al. 2011; Capizzi 2020). Depending on at what stage of eradication a program is stopped, the remaining population can rebuild to its pre-management densities or beyond (Armstrong et al. 2001; Bradshaw et al. 2007).

#### Approach

To examine how quickly the Kangaroo Island cat population would recover to initial population size ($N_1$ = 1629) relative to variation in the premature stopping of the culling program, we stopped the optimal management cull (0.5 proportionate annual cull) at yearly values following the initial cull (0.60 proportionate cull for first two years). We then allowed the population to rebuild using the same methods as in the untreated population (see main text *Methods*). We then recorded the time it took for the rebuilding population to equate to $N_1$. We ran each stoppage scenario for 10,000 iterations (randomly sampling 10,000 times from the stochastic survival and fertility vectors) to calculate the mean and 95% confidence limits for population size.

#### Results

Without a maintenance cull (i.e., only two years with 0.6 proportional cull applied), the population recovers to its initial size in 11 years (range: 7 – 32 years) (Fig. S4). Stopping culling prior to the end of the targeted duration of the 10-year culling program (i.e., 9$^{th}$ year) results in population recovery to initial size in 26 years (range: 17-58 years) (Fig. S4).

**Figure S4**. Years required for population to rebuild to founding population size ($N_1$) when the maintenance cull is stopped prematurely (year maintenance cull stopped). Initial culling of 0.60 for two years is followed by maintenance culling (0.5) for the years indicated. Solid black line represents the median time to rebuild to founding population (years), and the grey area represents the 95% confidence interval.

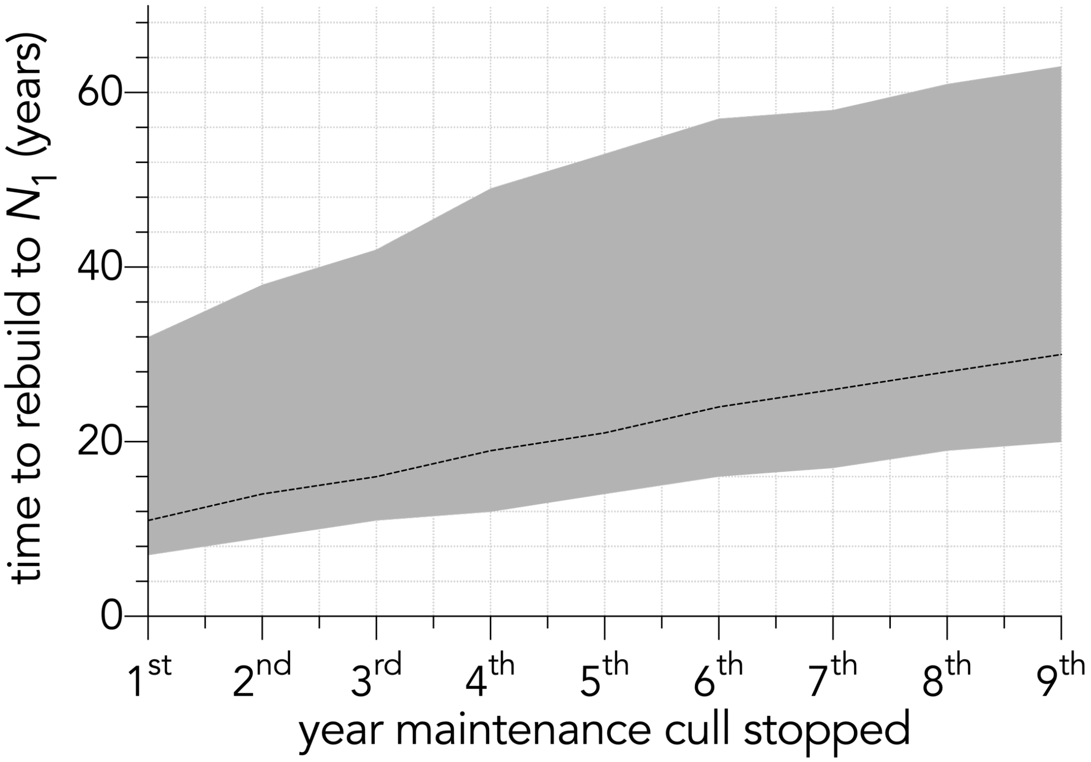

### Supporting information references

Armstrong DP, Perrott JK, Castro I. 2001. Estimating impacts of poison operations using mark-recapture analysis: hihi (Notiomystis cincta) on Mokoia Island. New Zealand Journal of Ecology:49-54.

Bradshaw CJA, Field IC, Bowman DMJS, Haynes C, Brook BW. 2007. Current and future threats from non-indigenous animal species in northern Australia: a spotlight on World Heritage Area Kakadu National Park. Wildlife Research **34**:419-436.

Capizzi D. 2020. A review of mammal eradications on Mediterranean islands. Mammal Review **50**:124-135.

Keitt B, Campbell K, Saunders A, Clout M, Wang Y, Heinz R, Newton K, Tershy B. 2011. The global islands invasive vertebrate eradication database: a tool to improve and facilitate restoration of island ecosystems. Island invasives: eradication and management. IUCN, Gland, Switzerland:74-77.

Natural Resource Kangaroo Island. 2015. Feral cat eradicaiton on Kangaroo Island 2015-2030 PROSPECTUS. Kangaroo Island.
